# Large language model inference of macromolecular complex composition via model consensus and experimental data integration

**DOI:** 10.64898/2026.05.20.726735

**Authors:** Mikhail Zhernevskii, Daniel Lynch, Vasilii Gorbunov, Yue Bao, Dmitry Korkin

## Abstract

Large language models (LLMs) are poised to reshape how biologists retrieve specialized knowledge at scale. Yet their performance on deep, domain-specific queries is poorly defined because much biological information resides in structured databases or large experimental datasets rather than in a free text format. One such gap in cellular biology lies in identifying major macromolecular complexes, conserved biological units essential to many cellular processes. Cataloging large complexes, such as the ribosome or RNA polymerase, along with their constituent genes, presents a significant challenge for LLMs because of their tendency to hallucinate and to produce incomplete or inconsistent lists of components. Here, we systematically evaluate six state-of-the-art LLMs on the task of retrieving the gene components of 91 protein complexes and develop an integrative framework that combines LLM output consensus with experimental multi-omics data to reconcile and filter model responses. We found that two extensions of a basic single-LLM baseline, (i) aggregating LLM outputs into a consensus and (ii) integrating LLM predictions with the experimental data, each improved retrieval accuracy. Furthermore, a consensus of LLM outputs integrated with the incomplete experimental data using a graph-theoretic approach achieved the highest accuracy (F1 score of 82.5%), compared to the best stand-alone singe LLM (F1 score of 76.4%). These results show that optimized integration of predictions from multiple LLMs and high-throughput experimental data can support scalable, semi-automated curation of specialized biological resources, providing a general template for benchmarking and deploying LLMs for the structured knowledge retrieval tasks in molecular biology.

## Main

Proteins in the cell rarely function in isolation. Instead, they form tightly connected macromolecular complexes, often in concert with RNA and DNA molecules, which underpin the majority of core cellular processes^1-3^. From ribosome to spliceosome, nuclear pore complex to RNA polymerase, there are hundreds of complexes, many consisting of dozens, even hundreds of proteins. Because of their intrinsic complexity, only a small fraction of complexes have been experimentally studied in detail. As a result, systematic information on macromolecular complexes remains limited, frequently lacking even fundamental details such as molecular composition, *i*.*e*., the list of genes encoding protein subunits comprising the complex. Large-scale experimental efforts often contain only partial information about the complexes. The Protein Data Bank^4^, a centralized resource on macromolecular structures, predominantly contains incomplete or fragmented complex structures, often missing multiple subunits. Similarly, large-scale high-throughput proteomics and interactomic efforts^5, 6^ provide valuable yet still partial datasets that cannot fully resolve the molecular composition of most complexes. Recently, efforts on creating a manually curated compendium of macromolecular complexes and their molecular compositions have been initiated^7^, but these efforts are far from being complete, as they are bound by time-intensive manual literature search and curation.

In the past few years, the information search and analysis tasks have been greatly advanced by the introduction of large language models (LLMs)^8-10^. LLMs are generative pre-trained transformers that leverage deep learning architectures and are trained on massive corpora of unlabeled text. Some of the prominent examples include ChatGPT, Claude, and Gemini^11, 12^. These models differ in their underlying architectures, training datasets, and degree of multimodal integration. Consequently, they may provide not only correct answers that vary from one another, but also make various kinds of mistakes. A common shortcoming that has been observed in the LLMs is their tendency to hallucinate. Hallucination, the property of LLMs to generate confident but factually incorrect or nonsensical outputs that are not grounded in real-world knowledge, is not merely an occasional mistake but an inherent feature of this technology, due to its generative nature^13, 14^. Hallucination-based errors are generally difficult to detect because the LLM outputs often appear coherent and contextually appropriate, making them indistinguishable from accurate information without external validation based on the domain-specific data. While the LLMs were originally designed to use in generic natural language processing tasks, they had been recently applied to address domain-specific questions, including those ones in molecular and cell biology. Specifically, two recent works have been introduced that address the important tasks of gene ontology (GO) functional annotation and cell type prediction^15, 16^. Both methods show promise in exploiting LLM-generated information; however, it is also evident from their assessment that an LLM alone has a long way to achieve accuracy comparable with the domain expert-based information retrieval, even for the reasonably straight-forward biological tasks.

In this Brief Communication, we explored the capacity of LLMs to extract the protein/gene composition of the major large (>10 distinct proteins with some exceptions, see below) macromolecular complexes. To do so, we addressed two related problems: Problem 1 was to retrieve a list of 100 names of such macromolecular complexes, and Problem 2 was to retrieve a list of genes whose protein products contribute to each complex. Furthermore, we asked if it was possible to improve the accuracy of the prediction by (1) developing a consensus-based approach that leveraged outputs from two or more LLMs, or (2) integrating LLM results with the high-throughput -omics data describing certain properties of these complexes. As a result, we developed and assessed four protocols that included six LLMs and three sources of the high-throughput experimental data: integrated proteomics data^5^, interactomics data^6^, and 3D molecular structures of protein complexes^4^. Four different information retrieval protocols were considered: (1) instructions to retrieve the specific information (referred to as ‘prompts’) submitted for individual LLM; (2) prompts for individual LLMs combined with the high-throughput experimental data; (3) consensus of LLM prompts; and (4) consensus of LLM prompts combined with the high-throughput experimental data (Methods, Fig. 2). Our assessment was done on a set of literature-curated human protein complexes, and on two sets of experimental high-throughput -omics datasets with the help of advanced graph-theoretic approaches to generate accuracy measures based on the incomplete experimental data (Methods).

We first assessed the performance of a single LLM, ChatGPT (GPT-4, version 4o), to determine biological names of macromolecular protein complexes with ten or more distinct genes contributing to the complex. We found that the LLM can accurately extract names of such protein complexes, retrieving 89 correct complexes out of the list of 100 (89% accuracy). The primary source of errors was identifying organelles and other large systems (*e*.*g*., Golgi apparatus and actin filament network) as macromolecular complexes (Suppl. Table S1). Another error was retrieval of complexes with less than 10 proteins: when compared the results of the LLM retrieval against a list of 28 verified complexes (Methods), we found that 11 complexes included between 5 and 8 unique proteins. Five complexes with the fewest number of five unique proteins retrieved by ChatGPT and verified during the curation process included DNA base excision repair complex, sister chromatid cohesion complex, condensins, replication factor C (RFC) complex, and synaptotagmin□SNARE complex.

Next, we evaluated the performances of six LLMs (Suppl. Table S2): ChatGPT (ver. o1-preview/o1), Claude (ver. 3.5 Sonnet), DeepSeek (ver. R1), Gemini (ver. 2.5 Pro), Llama (ver. 4), and Perplexity (using Llama 3), on the second Problem of predicting the gene components of macromolecular complexes, where for a curated set of 91 complexes (89 complexes validated from Problem 1, and two new complexes added to the list), each LLM was prompted to retrieve a set of unique genes whose protein products comprised the complex (Methods, Fig. 1A, Suppl. Fig S1). We first evaluated the performance over a set of 28 literature-curated well-studied macromolecular complexes (Fig. 1A). For all 28 complexes and specific LLM, the best-performing prompt according to the F1-score was selected (Suppl. Tables S2-S4, Suppl. Fig. S2, Methods). We found that overall 4-block prompt outperformed other prompts across different LLMs. We also found that the performance across different complexes and different LLMs varies significantly (Fig. 1A, Suppl. Fig S3). While for certain macromolecular assemblies, such as replication factor C complex or exocyst complex, all six LLMs report a perfect 100% F1 accuracy, for other complexes, such as helicase or histone deacetylase complex, several LLMs report poor performance, with only two LLMs (ChatGPT and Llama) and one LLM (Gemini) reaching the F1 accuracy greater than 60%, for the first and second complexes, respectively. Overall, ChatGPT outperformed other LLMs with the average F1 score of 76.3% over the 28 curated complexes.

**Figure 1.**
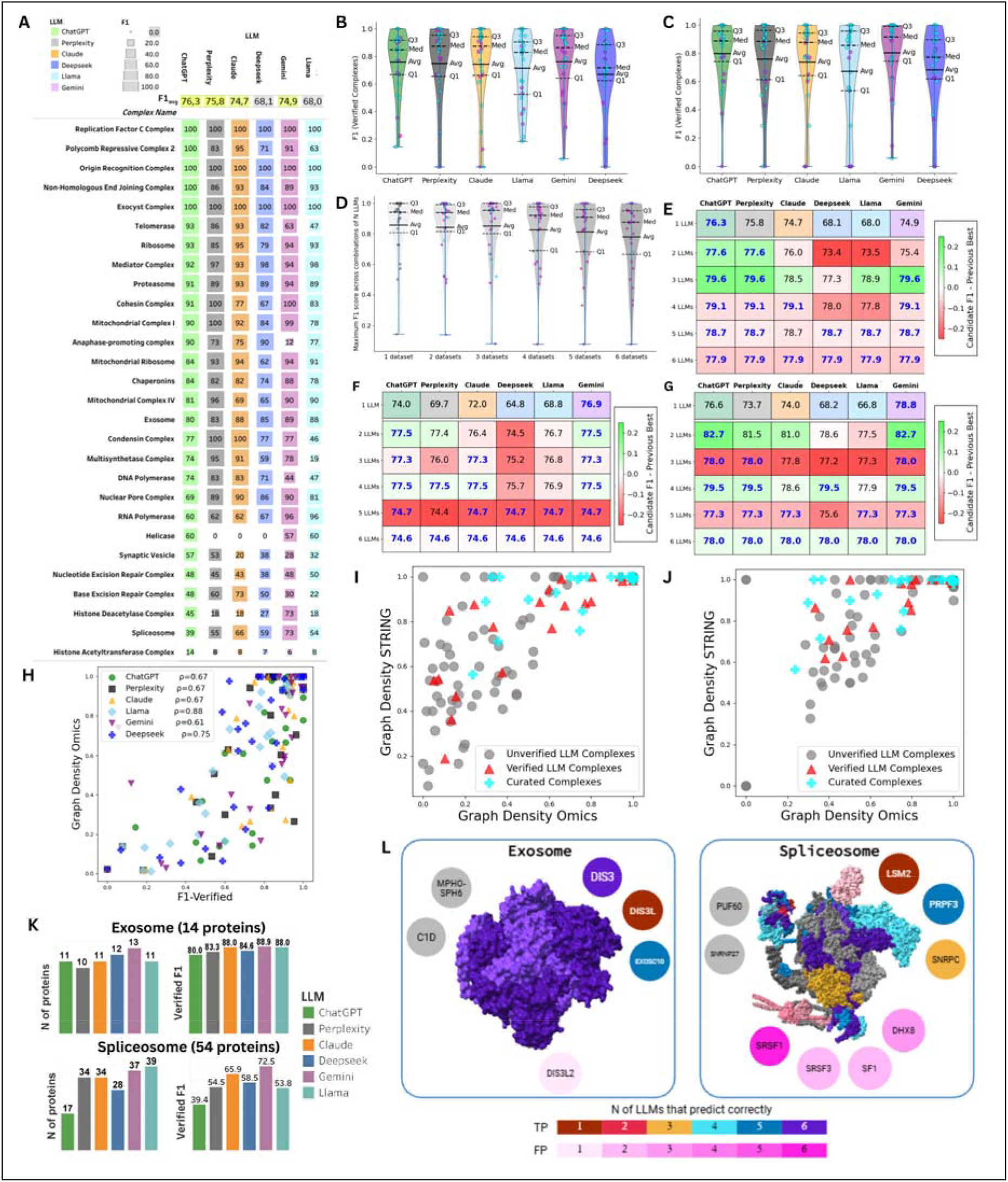
Performance of three strategies for predicting the protein composition of large protein complexes using six LLMs, with and without integration of high-throughput multi-omics data. **A:** Accuracy of retrieving components for 28 manually curated protein complexes, represented as an ‘F1-verified’ measure. Shown are the F1-verifeid scores for individual complexes and the averages across all 28 complexes calculated for each individual LLM. **B:** Changes in F1-verifeid prediction accuracy for LLMs’ output for the same dataset of 28 curated complexes after integrating predictions with experimental -omics data using the Binary Classification method. Cyan dots indicate an increase, and magenta dots indicate a decrease, in the F1-verified score for each complex after integration. **C:** Similar representation of F1-verified accuracy changes for the 28 curated complexes after integrating LLM outputs with the -omics data using the Bridges method. Cyan and magenta dots indicate the same changes as in **B. D:** Changes in the F1-verified accuracy of each complex between the most accurate single-LLM output (1 dataset, grey dots) and the most accurate majority-vote consensus of *N* LLMs, *N* = 2, …, 6. Cyan and magenta dots indicate the same changes as in **B. E:** The best average F1-verified accuracy calculated via the majority-vote consensus across all combinations of *N* LLMs, *N* = 1, 2, …, 6, that include a specific LLM (each column). Cell color for rows 2-6 represents the change between the accuracy of the current combination of *N* LLMs and the best accuracy among all possible combinations of (*N*-1) LLMs. **F:** Improvement in the retrieval accuracy (F1-verified score) by integrating the majority-vote consensus of *N* LLMs, *N* =1, 2, …, 6, with experimental -omics data using the Binary Classification method; cells are colored as in **E. G:** Improvement in the retrieval accuracy (F1-verified score) by integrating the majority-vote consensus of *N* LLMs, *N* =1, 2, …, 6, with experimental -omics data using the Bridges method; cells are colored as in **E. H:** Comparison of a Graph Density measure derived from -omics sources (hu.MAP, HuRI, PDB) with F1-verified accuracy measure for 28 curated protein complexes across six LLMs. Shown are the Spearman correlation coefficients, *ρ*, between the two measures. **I:** Assessment of 91 LLM complexes retrieved using the majority-vote consensus (based on the most accurate combination of two LLMs: ChatGPT, and Gemini) with two Graph Density measures: one using -omics sources and another one using STRING data. 28 verified LLM complexes are marked red. The corresponding manually curated complexes are marked as cyan. **J:** Similar to **I**, but retrieved by integrating the majority-vote consensus and experimental data using Bridges method. Integrating experimental data substantially improved both scores. Data points of 28 verified curated protein complexes remained unchanged (cyan). **K:** Case-studies of retrieving protein components for an ‘easy’ case (exosome, 14 protein components) and ‘difficult’ case (spliceosome, 54 protein components). For each LLM, the number of correctly predicted components and the corresponding F1-verified score are shown. **L:** Protein components of two structurally resolved complexes, exosome (PDB ID 2NN6) and spliceosome (PDB ID: 6FF7) predicted by individual LLMs. Each gene is represented either by a 3D structure (if structurally resolved) or by a circle (if unresolved). True Positive (TP) components, both structurally resolved and unresolved, are colored using a TP color scheme based on how many LLMs correctly predicted them. False Positive (FP) components are colored using a FP color scheme based on how many LLMs predicted them. False Negative (missed ground-truth components present in the curated complex but not predicted by any LLM) are colored grey.

**Figure 2.**
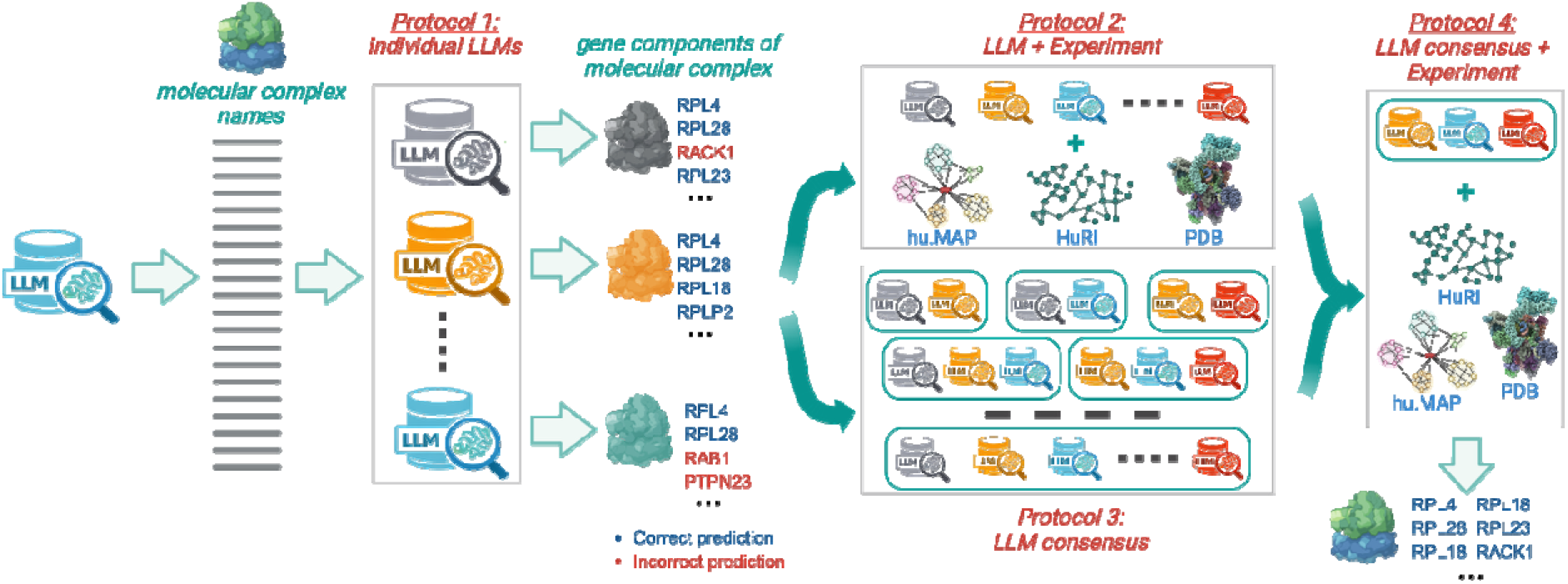
Overview of the four protocols to retrieve the gene content of a large macromolecular complex using six different LLMs and three large-scale sources of experimental information.

Manual, literature-based, curation of the complexes presents an accurate but time-consuming approach to assessment. To adopt a more high-throughput assessment protocol, we next quantified the consistency of the predicted complex composition with experimental large-scale -omics data. Specifically, we leveraged three different graph-driven measures (Methods): (1) *F1-huMAP*, where the positive and negative matches are defined by applying the Binary Classification Integration protocol to hu.MAP dataset; (2) *graph density (Omics)*, where the consistency of LLM predictions is assessed against all three large-scale experimental -omics sources for macromolecular complexes and protein-protein interactions, hu.MAP^5^, PDB^4^, and HuRI^6^, by constructing a macromolecular complex graph and measuring its density; and (3) *graph density (STRING)*, where the same is measured against STRING DB^17^, which includes both experimental and computational predictions on protein-protein interactions. Our analysis of correlations between the regular F1-verified score and the three experiment-based scores (Fig. 1H and Suppl. Fig. S4, S5, Suppl. Table S5) on the 28 verified complexes showed high correlation between the scores (Spearman correlation), suggesting that the latter scores could be adopted to assess LLM performance accuracies on larger datasets.

In addition to using the experimental -omics datasets for the assessment, these datasets could potentially be used to improve the accuracy of the LLM-based prediction through an integration of the experimental information sources. To this end, we next used the two graph-theoretic methods, Binary Classification and Bridges, for integrating incomplete large-scale omics information from the three sources described above (Figs. 1B, 1C, see *Methods* for more details). We found that while each of the omics sources provided partial information, leveraging these sources with the above two integration methods could improve the accuracy of LLM-based predictions, albeit not for all LLMs (Fig. 1B, 1C, Suppl. Tables S6, S7). The largest difference between the numbers of F1-improved and F1-worsened complexes was observed for Bridges integration with Gemini, with 39.3% of complexes improved and 21.4% worsened. Interestingly, for both integration methods, we observed that the probability of a score to be improved was higher for the complexes whose original F1 scores were above the average values. However overall, integrating experimental information using the Binary Classification method did not improve the original LLM accuracy for all but two LLMs (Suppl. Fig S6): Llama (improvement of average F1 score from 0.680 to 0.688) and Gemini (improvement from 0.749 to 0.769). Integrating experimental information using the Bridges method was slightly better, improving the accuracy for ChatGPT (from 0.763 to 0.766), DeepSeek (from 0.681 to 0.682), and Gemini (from 0.749 to 0.788), thus improving the best accuracy of an individual LLM from 0.763 (ChatGPT) to 0.788 (Gemini).

Next, we tested whether the accuracy of the information on protein complexes provided by individual LLMs can be improved by combining their predictions into a consensus of several LLMs. To determine how many and which LLMs constitute the most accurate consensus combination, we performed a comprehensive analysis of the consensus votes across all possible combinations of six LLMs (with consensus defined by a majority vote, *i*.*e*., at least 50% of all LLMs in a given combination). The analysis that covered these 57 (= 26 – 6 – 1) combinations, from all possible pairs of LLMs to all six LLMs, revealed that a consensus of three LLMs generally performed better than any other combination size (Fig. 1D, E). Furthermore, the combination of ChatGPT, Perplexity, and Gemini was the best among all other triplets, achieving F1 accuracy of ∼0.796 over the 28 curated complexes (Fig. 1E), again an improvement from the best individual LLM accuracy of 0.763 (ChatGPT).

Having separately evaluated LLM consensus and integrating LLM information with omics-based data, we next asked whether jointly integrating large-scale omics data with the LLM consensus could further improve the accuracy of determining the components of macromolecular complexes. Specifically, we used the same two -omics data integration methods, Binary Classification and Bridges, together with the 58 possible consensus combinations of LLMs. We found that integration via Binary Classification yielded only a marginal improvement over the consensus baseline, reaching F1 accuracy of ∼0.798 and achieving this accuracy on a slightly different consensus of 3 LLMs: ChatGPT, Perplexity, and Gemini. In contrast, Bridges Integration substantially improved the accuracy, reaching F1 score of ∼0.827. Surprisingly, the best consensus included not three but just two LLMs: ChatGPT and Gemini. In addition to improving accuracy for both above tasks, the Bridges -omics data integration method greatly improved the accuracy of the remaining complexes from the dataset of 91 complexes, as characterized by two measures, Graph Density Omics and Graph Density STRING (Fig. 1J), relative to the complexes and their components inferred by the individual LLMs (the best performing LLM are shown in Fig. 1I, Suppl. Fig. S8).

The biological domain-specific information retrieval tasks, like the one considered in this work, are a classic example of “hard to find, easy to verify” problem. Similar problems arise in cryptography (*e*.*g*., searching for a key versus verifying it), number theory (*e*.*g*., factoring an integer versus checking the product), or puzzle solving (*e*.*g*., finding versus validating a Sudoku solution). By analogy, the difficult task of finding the component of a protein complex can be delegated to an LLM, while verification remains with the human expert, either through targeted literature curation or by checking consistency with experimental data in biological databases. In our work, we show that information-retrieval capacity of LLMs for such tasks can be substantially improved by leveraging two basic principles: (1) aggregating LLM predictions into a multi-model consensus and (2) integrating LLM outputs with potentially incomplete omics data from large-scale datasets. At the same time, we observe clear limitations: for complexes that are poorly characterized in the literature or in experimental datasets, even consensus LLM predictions integrated with omics evidence can fail to recover the correct subunit composition. Some complexes are robustly retrieved across all LLMs, whereas others show highly variable and substantially lower accuracy (Fig. 1 K, L and Supp. Fig S1). A further challenge is posed by complexes in which only a subset of proteins form a permanent core, while additional proteins associate transiently during specific functional states. In this case, it would be critical for LLM-driven retrieval not only to identify the complex as a whole but also to distinguish between permanent and context-dependent components.

In conclusion, because current LLM models are trained primarily on text and rarely ingest raw domain-specific datasets directly, integrating LLM-based retrieval with curated, machine-readable biological resources offers a promising route toward a robust platform for biological knowledge extraction. Extending this strategy to catalogue macromolecular complexes across different species would be an important next step. Systematic evaluation of additional high-throughput modalities (*e*.*g*., co-localization, co-expression, genetic interaction) as evidence layers should further improve both the coverage and confidence of the approach. With LLMs growing more powerful, while still remaining diverse in terms of the outputs they produce and errors they make, the field of molecular biology is well positioned to harness them for information retrieval. And the field can best capitalize on this paradigm through principled data integration and carefully designed consensus strategies.

## Methods

### Method overview

We develop an integrative LLM-driven computational pipeline for retrieving large human macromolecular complexes (containing at least 5 distinct subunits) and their components (Fig. 1). We test two key hypotheses that the retrieval accuracy can be improved (1) by aggregating predictions across multiple LLMs into a consensus, and (2) by integrating LLM outputs with high-throughput experimental datasets. To evaluate these hypotheses, we incrementally refine the pipeline and assess performance at each stage using multiple accuracy metrics. We first compare the performance of six individual LLMs. Next, we integrate the predictions from each LLM with two types of high-throughput interactomic data: one derived from an integrated dataset containing large-scale yeast-two-hybrid (Y2H) and literature data on binary protein-protein interactions^6^, and another from tandem affinity purification mass spectroscopy (TAP-MS) data on the proteins participating in a complex^5^. We then systematically evaluate all possible consensus combinations of the six LLMs to determine whether the best-performing ensemble outperforms the best single model. Finally, we select the most accurate combination of LLMs and integrate its consensus predictions with the above experimental datasets. The performance of our approach is evaluated against a literature-curated reference set of protein complexes and partially available experimental data using both standard and novel graph-based metrics.

### LLM-based sources for macromolecular complex retrieval

We design LLM prompts around solving two main problems. For the first problem, we prompt ChatGPT (ver. 4o) to generate 100 human macromolecular complexes, with each complex requested in the prompt to include at least ten unique subunits. In our work, we focus on retrieving major macromolecular assemblies: large complexes that carry out central functions in the human cell and are likely to form predominantly permanent macromolecular interactions^1, 18, 19^. However, due to ‘hallucinations’ (outputs unfaithful to the source data, or unsupported by any known evidence or training context^13^), despite requesting only complexes with ten or more unique protein subunits in the LLM’s initial prompt, some of the retrieved complexes contained fewer than ten subunits. Therefore, for the second problem, the minimal requirement on the number of complex subunits is relaxed to five. The resulting list of complexes is then manually curated to remove erroneous, synonymous, or small complexes (<5 unique proteins), leading to the exclusion of 11 incorrect complex names (Suppl. Table S1). Furthermore, for the second problem, we augmented this LLM-derived set by adding two biologically important complexes that satisfied our original ≥10-subunit criterion but were not suggested by the LLM: the polycomb repressive complex 2 (PRC2) and the anaphase-promoting complex. Thus, for the second problem, the final working set comprises 91 macromolecular complexes (Supplementary Table S1). For each of these complexes, we then query each of the six LLMs to retrieve the full list of genes whose protein products are expected to participate in that complex.

The six LLMs-based methods used in this work include ChatGPT (ver. o1-preview/o1), Claude (ver. 3.5 Sonnet), DeepSeek (ver. R1), Gemini (ver. 2.5 Pro), Llama (ver. 4), and Perplexity (using Llama 3). The first five models all share the key LLM principles: they all have transformer-based architectures and are designed to carry out text-based natural language processing (NLP) tasks such as answering questions, summarizing text, and generating requested content. However, the models differ by their specific architectures and their complexities, training sets used, multimodal capabilities, and other features. The last model, Perplexity, employes Retrieval-Augmented Generation (RAG) architecture to integrate the information from several LLMs. Given the complexity of different scenarios and frequent updates of the models, for some LLMs, the data are generated using several consecutive model versions (Supplementary Table S2).

#### Prompt Engineering

The prompts are designed to be identical for all LLMs and aim at retrieving two types of information corresponding to the two tasks defined above (Supplementary Table S3): (1) the name of a human macromolecular complex, and (2) the complete list of human genes whose protein products contribute to that complex. Following curation of the final list of macromolecular complex names, we evaluated four prompting strategies for each LLM to maximize the accuracy and completeness of the retrieved gene lists. All LLMs are run using their default parameters. Given the large number of prompts evaluated, each prompt is issued only once. The four prompting strategies are as follows:

1. **“Zero-shot” prompting with encouragement**. In this strategy, the prompt does not include structured examples of the desired output, but instead provides only a protein name (*e*.*g*., ‘RPL23’) to ensure the proper output formatting. Encouragement is represented as a supplement to the main request to steer the model’s reasoning and emphasize task completion (Supplementary Table S4).
2. **“One-shot” prompting with encouragement**. In this strategy, the prompt includes a structured example of the desired output in which the macromolecular complex name matched to a list of constituent proteins. The generic structure of the output is provided as a part of the prompt in the following form (Supplementary Table S3): *{“example molecular complex”: [ “example protein 1”, “example protein 2”*…*]}* Encouragement is applied in the same manner as in the first strategy.
3. **Context prompting**. The prompt is organized in two parts. In the first part, we introduce the LLM to the structural context of our query: the format of the input (names of macromolecular complexes) and output (the list of proteins contributing to each complex), the desired processing mapping of the data, and the encouragement phrase. Then, in a separate message, we provide the LLM only with the list of protein complexes.
4. **Context and Splitting the Input**. This strategy extends Method 3 by partitioning the list of macromolecular complexes into *N* blocks of approximately 91/*N* complexes, with *N* = 2, 3, 4, 5, and 10 (rounded as needed, with the final block containing any remaining complexes). As a result, there are five prompt sequences in this prompt category. The goal of this strategy is to split the overall data retrieval task into smaller subtasks, thus asking to retrieve fewer proteins at once.

All prompting methods except Gemini are implemented using the web interface for ChatGPT, Claude, Perplexity, Llama, and DeepSeek. When generating the output, sometimes an LLM halts prematurely without providing information for the complete list of complexes. In this case, we resume generating the information by providing the LLM with a continuation cue:

“*Your list is not complete. Please continue generating proteins for the rest of the complexes in the list*.*”* If necessary, this strategy is applied repeatedly until the list of proteins is obtained for all 91 complexes. Followed by assessment of the individual prompting methods, the optimal prompt for each LLM is selected for downstream data integration analyses.

### Experimental sources for macromolecular complex information retrieval

To improve the accuracy of LLM-based information retrieval, we integrate LLM-generated predictions with information obtained from high-throughput experimental sources of protein-protein interactions (PPIs). Our primary source is the Human Protein Complex Map (hu.MAP)^5^. The hu.MAP dataset contains >7,000 protein complexes, and is constructed by integrating multiple large-scale experimental datasets, including affinity purification mass spectrometry (AP/MS) from Bioplex2.0^20^, biochemical fractionation^21^, proximity labeling, and RNA hairpin pulldown experiments^22^. We download the protein complexes file from the hu.MAP online repository (https://humap2.proteincomplexes.org/static/downloads/humap2/humap2_complexes_20200809.txt) and repository and convert it into two JSON files for downstream use: the first file contains complexes as keys and associated proteins as values, while the second file contains proteins as keys and complexes that contain those proteins as values.

As the second source of interaction data, we use the Human Reference Interactome (HuRI)^6^, a systematic proteome-wide reference map of the PPI network (interactome). The current HuRI release contains >64,000 PPIs mediated by >9,000 human proteins derived from several yeast two-hybrid (Y2H) assays. Although incomplete, HuRI provides high-confidence support for binary PPIs. We download and process the HuRI dataset into a JSON file, in which each gene is a key and its experimentally supported interaction partners are stored as values.

The last experimental source of the macromolecular complexes and their compositions used in this work is the Protein Data Bank (PDB), a comprehensive resource of structurally resolved macromolecules^4^. While covering the full range of biomolecules, PDB also includes structurally resolved macromolecular complexes. For each seed protein, we queried the PDB API to retrieve all associated macromolecular assemblies. For each structure, we extracted the list of protein components and mapped them to Ensembl gene identifiers. This procedure yields, for each protein appearing in an LLM-predicted complex, the set of PDB structures that contain it, and for each structure, the list of proteins that comprise the complex. These data form the basis of PPI graphs used in graph-based analyses described in the next section.

### Integrating LLM and experimental interactomics data

We next assessed whether predictions from individual LLMs can be improved by integrating them with the high-throughput experimental data. Specifically, to integrate the protein complex predictions generated by LLMs with the data extracted from the three experimental sources, hu.MAP ^5^, HuRI ^6^, and PDB ^4^ databases, we developed two integration protocols: (1) Binary Classification Integration and (2) Bridges Integration.

#### Binary Classification Integration

For the first protocol, we use the data from a single experimental source, hu.MAP, which provides the broadest complex coverage among the three datasets considered. For each LLM complex, we first identify all hu.MAP entries (*i*.*e*., complexes) that share at least two proteins with that LLM complex. Second, within each of these hu.MAP complexes, we treat all constituent proteins as mutually interacting (*i*.*e*., we assume a fully connected complex and generate all possible interaction pairs). We then use the hu.MAP-defined interaction information to label proteins associated with the LLM complex. Each protein in the LLM complex is classified as:

1. True Positive (TP), if it appears in the LLM complex and has at least one hu.MAP-defined interaction with another protein from the same LLM complex.
2. False Positive (FP), if it appears in the LLM complex, but has no hu.MAP-defined interaction with any other protein from that LLM complex.
3. False Negative (FN), if it is not present in the LLM complex but is found in at least six of the above-identified hu.MAP complexes (*i*.*e*., complexes that share at least two proteins with the LLM complex).

Once all proteins are labeled, we construct an integrated complex for each LLM prediction by (1) retaining all TP proteins, (2) removing all FP proteins, and (3) adding all FN proteins to the LLM-derived complex.

#### Bridges Integration

The Bridges Integration protocol leverages experimental data from all three experimental sources (hu.MAP, HURI, and PDB) using graph-theoretic properties of a graph formed by the experimentally validated PPIs. For each LLM complex, we define a macromolecular complex graph whose nodes correspond to proteins in the complex and whose edges correspond to experimentally supported PPIs. While nodes can be directly obtained from the LLM complex prediction, the edges cannot. The tentative edges of the graph are therefore inferred from hu.MAP, HuRI, and PDB data.

First, for each LLM complex, we query hu.MAP and extract all complexes that contain at least two proteins from the LLM complex. This can yield multiple hu.MAP complexes for the same LLM complex. For each such hu.MAP complex, we restrict the protein set to those shared with the LLM complex and add the corresponding nodes to the macromolecular complex graph, if they are not present in the graph. We then connect all pairs of these nodes with edges, forming a fully connected subgraph (clique). This definition is intended to capture all potential interactions between the proteins within a hu.MAP-defined complex. Not that when two or more hu.MAP complexes overlap with the same LLM complex, there will be two or more cliques generated.

We apply an analogous procedure to PDB-derived assemblies that share at least two proteins with the LLM complex: we keep only proteins shared with the LLM complex and connect all corresponding node pairs with edges, again forming cliques. Lastly, because HuRI reports pairwise PPIs rather than complexes, we select HuRI interactions where both partners belong to the same LLM complex. To do so, for each protein in an LLM complex, we extract its Ensembl ID and use it to determine the list of all pairs of the proteins from the LLM complex in HuRI-sourced JSON file described above.

We then merge the PPI information from the three sources into a single integrated macromolecular complex graph. Specifically, the node set consists of all proteins in the LLM complex; an edge is added between a pair of nodes if that interaction is supported by any of the three source-specific macromolecular complex graphs. For each integrated graph, we then apply the graph-theoretic concept of 2-edge-connectivity. A connected graph is 2-edge-connected if it remains connected after the removal of any single edge. The intuition is that a biologically coherent macromolecular complex is unlikely to fragment into disconnected parts when a single interaction is lost. To identify 2-edge-connected components in the integrated graph, we first detect all bridges. A bridge is defined as a graph edge whose removal increases the graph’s number of connected components (where a component can be as small as a single node). We then remove all bridges in a graph, resulting in one or more 2-edge-connected components. The largest 2-edge-connected component is identified as the final molecular complex graph. In the rare case where multiple components have the same maximal size, one is selected at random.

### LLM output consensus voting strategies, with and without experimental integration

To determine if combining outputs from multiple LLMs outperforms any single model, we develop and evaluate a series of consensus protocols. For each of the 91 protein complexes, we considered all non-trivial combinations of the six LLMs, starting from all possible pairs and proceeding through triplets, quadruplets and quintuplets up to the full set of six models (2^6^ – 6 – 1= 57 unique combinations of the six LLMs). For each LLM subset, we constructed a majority-vote consensus: a protein was retained in the consensus complex if it was predicted by at least half of the LLMs in that subset.

As a final strategy, we combine majority-vote consensus predictions with the two experimental integration protocols described above, Binary Classification Integration and Bridges Integration. Specifically, for consensus combined with Binary Classification Integration, a protein is included in the final complex only if it (1) is predicted by at least half of the LLMs in the subset (majority vote) and (2) is retained in the complex reconstructed by the Binary Classification Integration protocol (see above). Similarly, for consensus combined with Bridges Integration, a protein is included only if it (1) passes the majority-vote criterion and (2) belongs to the experimentally supported complex derived from the Bridges Integration protocol.

### A curated dataset of macromolecular complexes and their components

All LLM-based protocols are initially evaluated on a “golden set” of 28 verified macromolecular complexes extracted from the CORUM database^7^ and further curated. CORUM is a manually annotated resource of mammalian protein complexes, providing information on complex function, subcellular localization, subunit composition, literature references, and other descriptors. Its annotations are derived from individual experiments reported in the peer-reviewed literature, with data from high-throughput screens explicitly excluded. To construct our golden set, we start from the previously retrieved list of 91 macromolecular complex names and queried CORUM (version 5.0) for complexes with identical or closely matching names, extracting a JSON file containing human macromolecular complexes and their constituent genes. Then, a final list of 28 complexes is selected and manually curated, requiring that each macromolecular assembly be supported by primary peer-reviewed publications. For three complexes (condensin, DNA polymerase and the nucleotide excision repair complex) CORUM contains multiple related entries; in these cases, we merge the corresponding subunit lists into a single composite complex definition. We refer to this curated set of 28 assemblies as the *verified complexes* and use it as the primary reference for benchmarking our LLM-based retrieval and integration strategies.

### Evaluation of LLM-based information retrieval strategies

Two major LLM prediction tasks are evaluated: (1) accurate prediction of protein names, *i*.*e*., whether a predicted human protein name corresponds to a real protein rather than a hallucinated entity; and (2) accurate prediction of the protein composition for a macromolecular complex, *i*.*e*., whether the set of subunits assigned to a complex matches reference data. For the first task, we quantify hallucination using a hallucination rate, as the number of hallucinated protein names divided by the total number of proteins in the LLM complex. For the second task, we compare LLM-predicted complex compositions against experimental sources as well as against the verified complex set, using four complementary metrics: (1) F1-Verified, (2) F1-HuMAP, (3) graph density based on -omics sources, and (4) graph density based on the STRING database. Each measure is described in detail below.

To compute the hallucination rate, each predicted protein name is validated against the human proteome using two centralized resources, Ensembl^23^ and STRING^17^, accessed via their APIs. A protein name is considered genuine if it is present in the human proteome of at least one of these databases; otherwise, it is classified as hallucinated.

For the second task, before we assess the accuracy of the LLM-generated predictions against verified data, we first evaluate the consistency of predictions across the six LLMs by comparing them to each other. For each pair of LLMs, we compare their predicted protein lists for each of the 91 macromolecular complexes using the F1 measure and then averaged across complexes. For a given LLM pair, the average F1-LLM score is defined as:

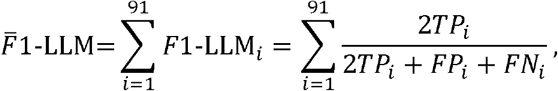

where true positives for complex *i* (*TP*_*i*_)represent the number of proteins shared by both LLM predictions, false positives (*FP*_*i*_) represent the number of proteins in the first LLM prediction not found in the second LLM prediction, and false negatives (*FN*_*i*_) represent the number of proteins in the second LLM prediction not found in the first one.

#### F1-Verified score

We first evaluate LLM-predicted complexes and their compositions by comparing them directly to the verified complexes using an F1 measure, which we referred to as F1-Verified score. While this benchmark set has limited coverage, it provides the most stringent and accurate assessment of LLM-based predictions. For each LLM-predicted complex that can be paired with a verified complex, we compare their compositions in terms of the genes whose protein products are assigned to that complex. The F1-Verified score is then calculated as:

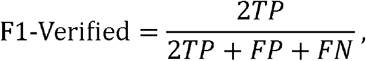

where true positives (*TP*) are the number of proteins shared between the LLM-predicted complex and verified complex, false positives (*FP*) are proteins present only in the LLM-predicted complex and not found in the verified complex, and false negatives (*FN*) are proteins present only in the verified complex.

#### F1-huMAP score

We next evaluate the LLM-predicted complexes against the high-throughput experimental complex dataset hu.MAP. This dataset offers substantially broader coverage than the verified list, but its annotations are not always complete, e.g., experimental evidence may exist for subcomplexes rather than for an entire macromolecular assembly. To compare LLM predictions against the hu.MAP data, we first define the positive and negative matches by reusing the Binary Classification protocol (see above). Specifically, for each LLM-predicted complex, we identify hu.MAP-supported true positives (*TP*_*1*_), false positives (*FP*_*1*_), and false negatives (*FN*_*1*_). Then, the following F1 score is calculated:

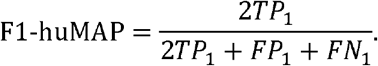

#### Graph Density (Omics Sources)

We next use a graph-theoretic measure to assess how strongly LLM-predicted complexes are supported by all three large-scale experimental sources for macromolecular complexes and PPIs: hu.MAP, PDB, and HuRI. For each LLM-derived complex, we first extract the relevant information from each source and construct a macromolecular complex graph following the procedure previously described for the Bridges Integration protocol (see above). The graph building procedure then yields: (1) one or more (when multiple hu.MAP complexes or subcomplexes map to the same LLM complex) fully connected graphs from hu.MAP data; (2) one or more (when multiple PDB structures correspond to the same complex) fully connected graphs from PDB data; and (3) a set of binary interaction edges from HuRI. We then integrate proteins and interactions from all three sources into a single macromolecular complex graph by merging edge information. The resulting integrated graph contains nodes corresponding to all proteins in the LLM-predicted complex, and an edge is placed between two nodes if that edge exists in at least one of the three source-specific graphs. To quantify the degree of experimental support for an LLM-predicted complex, we define a graph density measure:

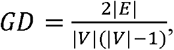

where |*V*| is the number of nodes in the integrated macromolecular complex graph (*i*.*e*., the number of proteins in the LLM-predicted complex) and |*E*| is the number of edges in the graph. Thus, the formula is the ratio of the number of potential PPIs supported by the experimental sources and the number of all possible interactions a macromolecular complex of |*V*| proteins can have.

#### Graph Density (STRING)

Finally, we assess the consistency of the LLM predictions with an additional PPI resource, STRING database^17^. Unlike the previous three sources hu.MAP, PDB and HuRI, which are based on the experimental data, STRING also contains predicted interactions and therefore is assessed separately. For each LLM-predicted complex, we collect STRING interactions among its proteins in the same manner as for HuRI, but restrict ourselves to high-confidence interactions (STRING confidence score > 0.8), retrieved via the STRING API. All such interactions among proteins in a given LLM complex define a STRING-based complex graph. We then compute graph density for this graph in the same way as before:

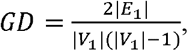

where |*V*_1_| is the number of nodes in the STRING-based complex graph (equal to the number of proteins in the LLM-predicted complex) and |*E*_1_| is the number of edges in that graph.

## Data Availability

All data used in this paper are publicly available. The hu.MAP dataset is downloaded from http://humap2.proteincomplexes.org. The HuRI dataset is downloaded from http://interactome-atlas.org. The CORUM dataset is downloaded from https://mips.helmholtz-muenchen.de/corum. StringDB is accessed through API (<u>string-db.org/api/json/get_string_ids</u>). PDB data is accessed through API (https://search.rcsb.org/redoc/index.html).

## Code Availability

The pipeline of data acquiring, processing, integration, and evaluation is available here at https://github.com/KorkinLab/LLM_Complexes/$repo

## Ethics Declaration

The authors declare no competing interests.

