## Supplementary Materials for "Large language model inference of macromolecular complex composition via model consensus and experimental data integration"

**Supplementary Information**

**Supplementary Tables**

**Supplementary Table S1.** List of 102 complexes, including (1) complexes correctly predicted by the LLM, with a manually verified list of gene components; (2) complexes correctly predicted by the LLM, without a manually verified list of gene components; (3) complexes manually added to the verified set; and (4) complexes incorrectly predicted by the LLM, together with the reasons for their exclusion. For each of the six LLMs, the XXX calculated for each complex is included.

**Supplementary Table S2.** Overview of selected LLMs used in this work for Problems 1 and 2
(* indicates an estimated number).

| **LLM** | **Version** | **Model size** | **Context length** | **Developer** |
| --- | --- | --- | --- | --- |
| ChatGPT | o1 | 300B* | 128K | OpenAI |
|  | 4o | 200B* | 128K |  |
| Claude | 3.5 Sonnet | 175B | 200K | Anthropic |
| Llama | 4 | 405B | 128K | Meta AI |
| Gemini | 2.5 Pro | 200B* | 128K | Google DeepMind |
| DeepSeek | R1 | 671B | 128K | Hangzhou DeepSeek AI |
| Perplexity | uses Llama3 | N/A | N/A | Perplexity AI |

**Supplementary Table S3.** Summary of LLM prompts. Asterisks mark the most accurate prompts for the evaluation of Problem 2 to predict gene components of a macromolecular complex.

| **LLM** | **Prompt Type** | | | |
| --- | --- | --- | --- | --- |
|  | **Zero-shot with encouragement** | **Context & Blocks** | **Context** | **One-Shot** |
| ChatGPT | Yes* | 2, 3, 4, 5, 10 | Yes | Yes |
| Perplexity | Yes | 2, 3*, 4, 5, 10 | Yes | Yes |
| Claude | Yes | 2, 3, 4*, 5, 10 | Yes | Yes |
| Gemini | Yes | 2, 3*, 4, 5, 10 | Yes | Yes |
| Llama | Yes | 2, 3, 4*, 5, 10 | Yes | Yes |
| DeepSeek | Yes | 2, 3, 4*, 5, 10 | Yes | Yes |

**Supplementary Table S4.** Description of the prompt formats provided for each LLM.

| **Prompt** | **Message** | **Additional Prompt** |
| --- | --- | --- |
| Zero shot | Please give me the list of all proteins that constitute the following human macromolecular complexes: [list of 100 macromolecular complexes]. Give me a JSON file with keys that are the human macromolecular complexes and all its values should be the IDs of the proteins that constitute them. Please use HGNC formatting. Do not have subkeys for any macromolecular complexes, just put all proteins in one value for each key. I do not care how complicated this task is. I believe in you and just want your best answer. It does not have to be completely accurate. I don't care how long it takes. Use any resources necessary to complete this task. I will not accept no for an answer. |  |
| One Shot | Please give me the list of all proteins that constitute the following human macromolecular complexes: [list of 100 macromolecular complexes].  Give me a JSON file with keys that are the human macromolecular complexes and all its values should be the IDs of the proteins that constitute them. Please use HGNC formatting (*e.g.* RPL23). Do not have subkeys for any macromolecular complexes, just put all proteins in one value for each key. I do not care how complicated this task is. I believe in you and just want your best answer. It does not have to be completely accurate. I don't care how long it takes. Use any resources necessary to complete this task. I will not accept no for an answer. Here is an example of a JSON formatted in the way I would like it to be when you produce one: { "example molecular machine": [ "example protein 1", "example protein 2"... ] } but you should include every molecular machine on my list of macromolecular complexes and include every protein that constitutes it. |  |
| Context | I am about to ask you to retrieve data about proteins that constitute certain human macromolecular complexes. You do not have to give me any information just yet, I am just preparing you and putting you in the right mindset. You will have to provide me with a JSON file with keys that are the human macromolecular complexes and all its values should be the IDs of the proteins that constitute them. Make your answer as comprehensive as you can. Please use HGNC formatting (*e.g.* RPL23). I do not care how complicated this task is. I believe in you and just want your best answer. I don't care how long it takes. Use any resources necessary to complete this task. I will not accept no for an answer. | Please give me the list of all proteins that constitute the following 100 human macromolecular complexes as JSON: [list of 91 macromolecular complexes]. |
| Context & 2 Blocks | The next 2 messages will each be a list with about 50 macromolecular complexes, accumulating to 100 total. You do not have to give me any information just yet, I am just preparing you and putting you in the right mindset. For each of them you will have to provide the list of proteins comprising them. You will have to provide me with a JSON file with keys that are the human macromolecular complexes matched to the list of IDs of the proteins that constitute them. Make your answer as comprehensive as you can. Please use HGNC formatting (*e.g.* RPL23). I believe in you and just want your best answer. I will not accept no for an answer. |  |
| Context & 3 Blocks | The next 3 messages will each be a list with about 33 macromolecular complexes, accumulating to 100 total. You do not have to give me any information just yet, I am just preparing you and putting you in the right mindset. For each of them you will have to provide the list of proteins comprising them. You will have to provide me with a JSON file with keys that are the human macromolecular complexes matched to the list of IDs of the proteins that constitute them. Make your answer as comprehensive as you can. Please use HGNC formatting (*e.g.* RPL23). I believe in you and just want your best answer. I will not accept no for an answer. |  |
| Context & 4 Blocks | The next 4 messages will each be a list with about 25 macromolecular complexes, accumulating to 100 total. You do not have to give me any information just yet, I am just preparing you and putting you in the right mindset. For each of them you will have to provide the list of proteins comprising them. You will have to provide me with a JSON file with keys that are the human macromolecular complexes matched to the list of IDs of the proteins that constitute them. Make your answer as comprehensive as you can. Please use HGNC formatting (*e.g.* RPL23). I believe in you and just want your best answer. I will not accept no for an answer. |  |
| Context & 5 Blocks | The next 5 messages will each be a list with about 20 macromolecular complexes, accumulating to 100 total. You do not have to give me any information just yet, I am just preparing you and putting you in the right mindset. For each of them you will have to provide the list of proteins comprising them. You will have to provide me with a JSON file with keys that are the human macromolecular complexes matched to the list of IDs of the proteins that constitute them. Make your answer as comprehensive as you can. Please use HGNC formatting (*e.g.* RPL23). I believe in you and just want your best answer. I will not accept no for an answer. |  |
| Context & 10 Blocks | The next 10 messages will each be a list with about 10 macromolecular complexes, accumulating to 10 total. You do not have to give me any information just yet, I am just preparing you and putting you in the right mindset. For each of them you will have to provide the list of proteins comprising them. You will have to provide me with a JSON file with keys that are the human macromolecular complexes matched to the list of IDs of the proteins that constitute them. Make your answer as comprehensive as you can. Please use HGNC formatting (*e.g.* RPL23). I believe in you and just want your best answer. I will not accept no for an answer. |  |

**Supplementary Table S5.** Comparison of a Graph Density measure derived from -omics sources with F1-verified accuracy measure for 28 curated protein complexes across six LLMs. Presented are the Spearman correlation coefficients, *ρ*, between the two measures and the corresponding P-values.

| **Model** | **Spearman ρ** | ***P*‑value** |
| --- | --- | --- |
| ChatGPT ● | 0.67 | < 0.0001 |
| Perplexity ■ | 0.67 | < 0.0001 |
| Claude ▲ | 0.67 | 0.0001 |
| Llama  ◆ | 0.88 | < 0.0001 |
| Gemini ▼ | 0.61 | 0.0005 |
| Deepseek **✙** | 0.75 | < 0.0001 |

**Supplementary Table S6.** Changes in F1-verifeid prediction accuracy for LLMs' output for the dataset of 28 curated complexes after integrating predictions with experimental -omics data using the Binary Classification method.

| **LLM/Change** | **Better** | **Worse** | **Neutral** |
| --- | --- | --- | --- |
| ChatGPT | 8 | 13 | 7 |
| Perplexity | 8 | 11 | 9 |
| Claude | 9 | 10 | 9 |
| Llama | 12 | 9 | 7 |
| Gemini | 10 | 9 | 9 |
| Deepseek | 6 | 13 | 9 |

**Supplementary Table S7.** Changes in F1-verifeid prediction accuracy for LLMs' output for the dataset of 28 curated complexes after integrating predictions with experimental -omics data using the Bridges method.

| **LLM/Change** | **Better** | **Worse** | **Neutral** |
| --- | --- | --- | --- |
| ChatGPT | 9 | 10 | 9 |
| Perplexity | 9 | 8 | 11 |
| Claude | 7 | 9 | 12 |
| Llama | 11 | 9 | 8 |
| Gemini | 11 | 6 | 11 |
| Deepseek | 10 | 8 | 10 |

**Supplementary Table S8.** Description of outlier complexes from Figure 1I.

| **Complex name** | **Graph Density**  **(Omics)** | **Graph Density**  **(STRING)** |
| --- | --- | --- |
| Cholesterol Biosynthesis Complex | 0.000 | 0.000 |
| Endoplasmic Reticulum-Associated  Degradation Complex | 0.000 | 0.000 |
| Glutamate Decarboxylase Complex | 0.000 | 0.000 |
| Protease Inhibitor Complex | 0.000 | 0.000 |
| RNA Editing Complex | 0.000 | 0.000 |
| Ribosome-Associated Quality Control Complex | 0.000 | 0.000 |
| Transporter Protein Complex | 0.000 | 0.000 |
| ER-Mitochondria Contact Site Complex | 0.000 | 1.000 |
| Membrane Type 1 Matrix Metalloproteinase Complex | 0.000 | 1.000 |

**Supplementary Figures**

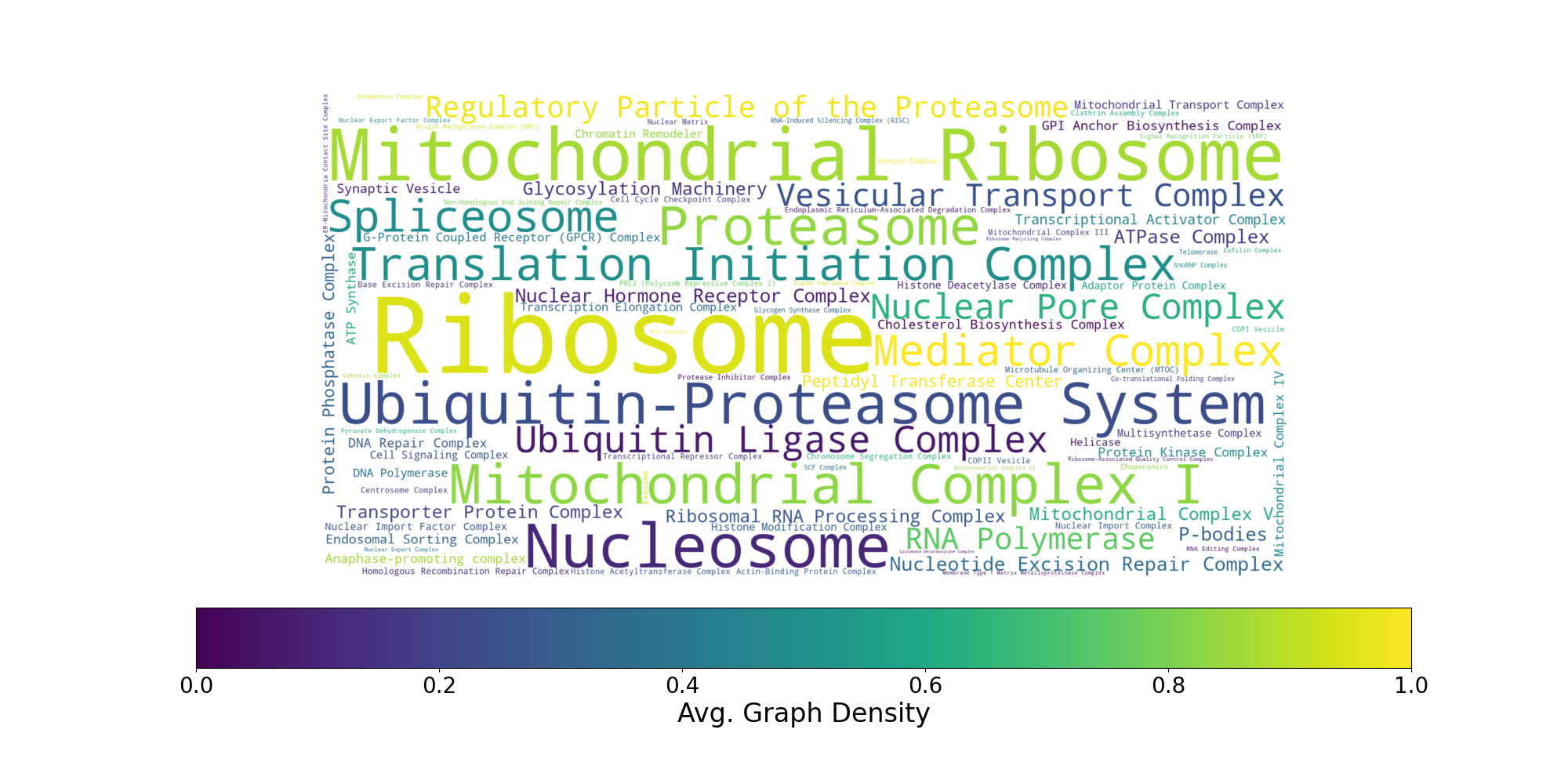

**Supplementary Figure S1.** Word cloud representation of the curated set of 91 macromolecular complexes. The size of each word is the average number of gene components predicted across 6 LLMs. The color (red, yellow, green) ranges based on average Graph Density across 6 LLMs.

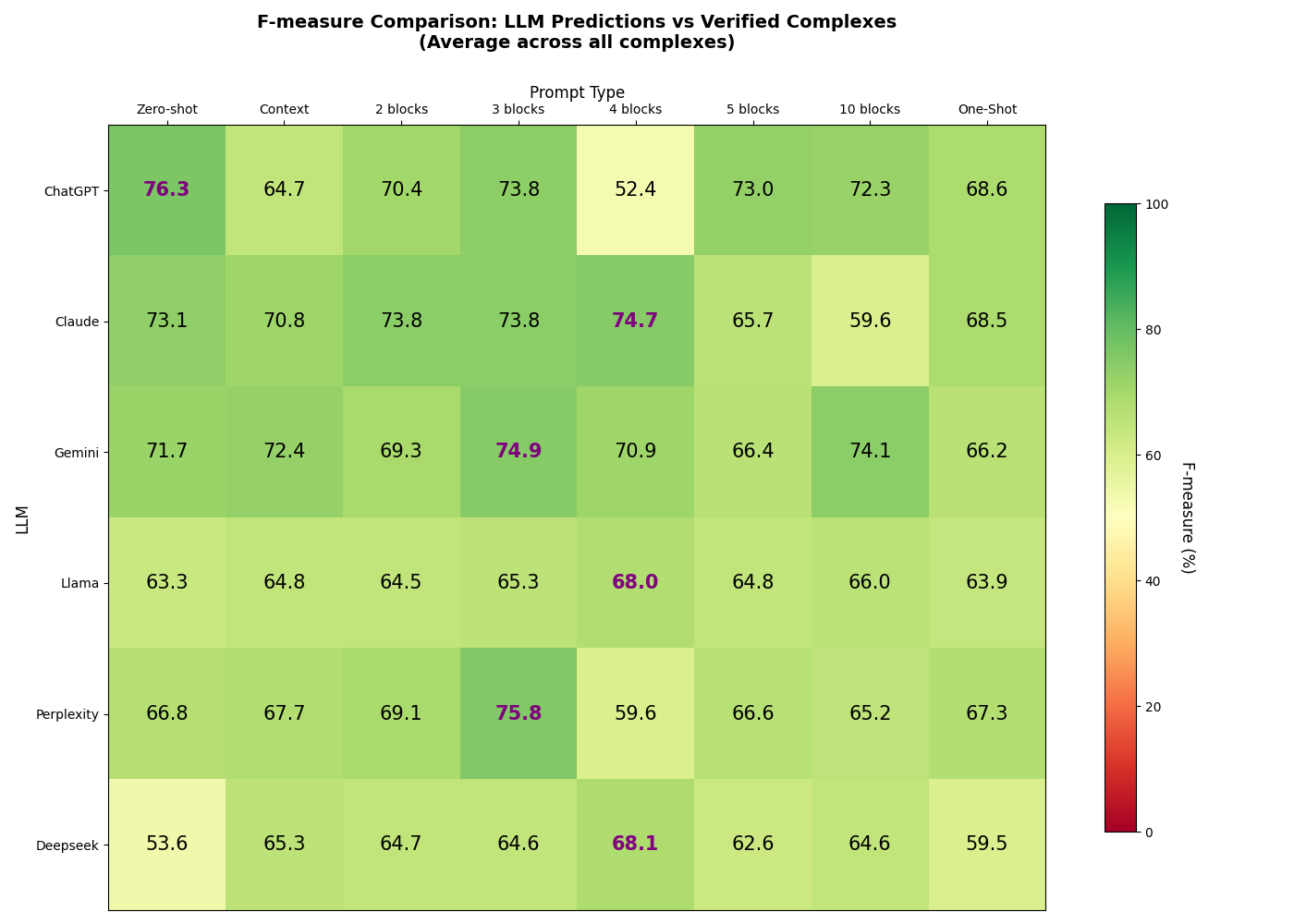

**Supplementary Figure S2.** The heatmap of F1 scores of individual LLMs across various prompting strategies for predicting gene content of 28 literature-curated macromolecular complexes (Problem 2). The highest-scoring prompt for each LLM is highlighted in bold purple and is selected as the final prediction for each LLM.

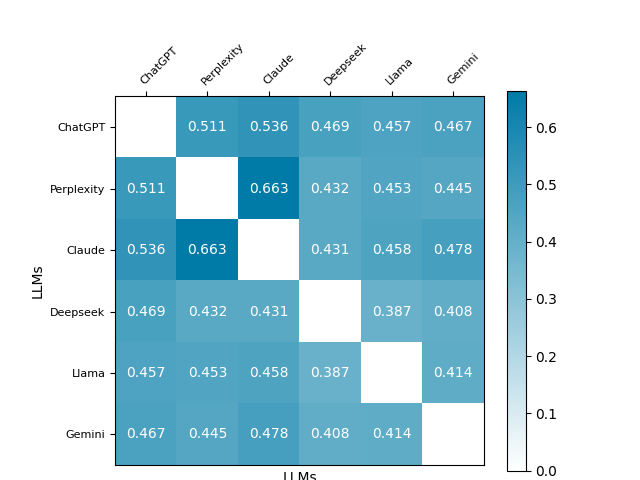

**Supplementary Figure S3.** The similarity between the outputs of each LLM calculated as a Tanimoto coefficient: *T_ij_=S_i_*∩*S_j_*/*S_i_*∪*S_j_*

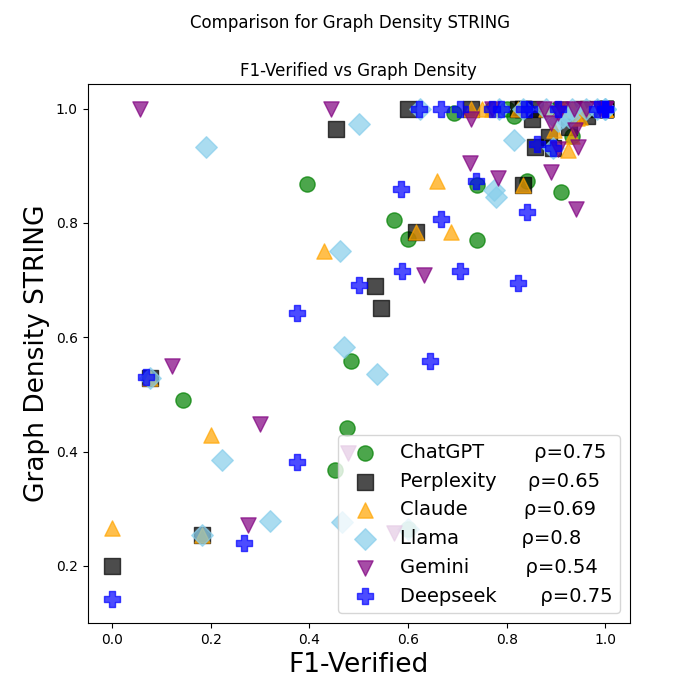

**Supplementary Figure S4.** Comparison of a Graph Density measure derived from STRING with F1-verified accuracy measure for 28 curated protein complexes across six LLMs. Shown are the Spearman correlation coefficients, *ρ*, between the two measures.

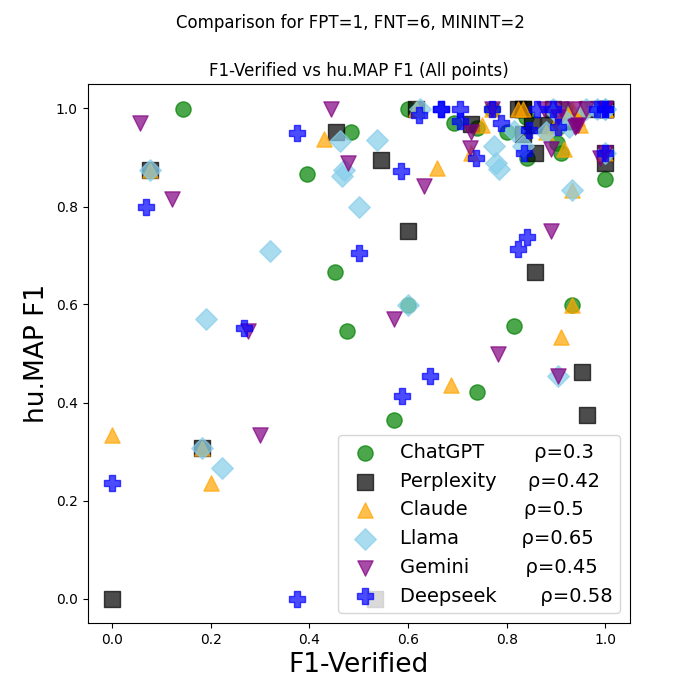

**Supplementary Figure S5.** Comparison of hu.mAP-based F1 score to the verified curated-complexes-based F1 among 28 protein complexes, grouped by LLM.

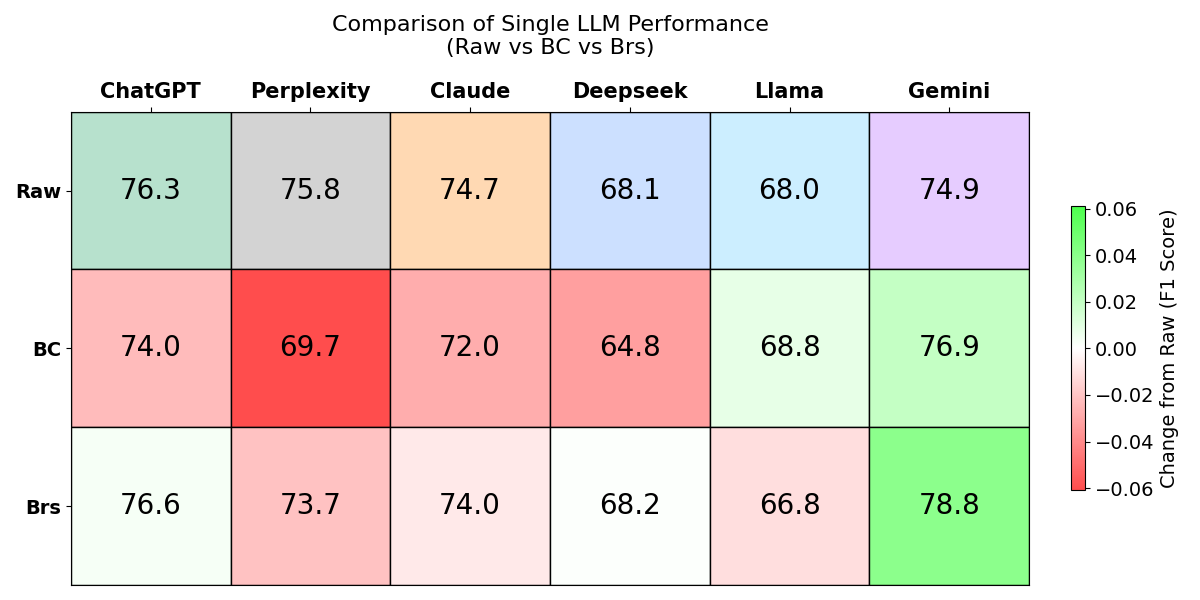

**Supplementary Figure S6.** This heatmap compares the F1 scores (percent) of individual LLMs across two experimental data integration methods, Binary Classification (BC) and Bridges (Brs), where the first row shows the baseline performance and subsequent rows illustrate the relative improvement or decline introduced by the Binary Classification and Bridges strategies.
